# Volatile cues from pathogenic, mutualistic and saprotrophic fungi cause specific, fungus-dependent responses in poplar

**DOI:** 10.1101/2025.10.07.680861

**Authors:** Peiyuan Zhu, Baris Weber, Maaria Rosenkranz, Andrea Ghirardo, Jörg-Peter Schnitzler

## Abstract

Plants are exposed to complex interactions with belowground organisms, yet how they differentiate between mutualistic and pathogenic fungi before physical contact remains largely unknown. We exposed the roots of young *Populus × canescens* to volatile organic compounds (VOCs) emitted by either a pathogenic (*Heterobasidion annosum*), a saprotrophic (*Postia placenta*), or an ectomycorrhizal (*Laccaria bicolor*) fungus. VOC analysis of the shared rhizosphere headspace and leaf emissions revealed that poplar plants could perceive and respond to fungal identity solely through airborne cues. The root-zone headspace contained fungus-specific sesquiterpene fingerprints that remained similar after three and six weeks of co-cultivation: Pathogen-derived VOCs induced constant high sesquiterpene emissions from the root-zone, whereas mycorrhiza caused low but targeted emissions of specific sesquiterpenes. In contrast, saprotrophic VOCs caused a temporal shift in root-zone VOC pattern, with increased sesquiterpene emissions after six weeks. Fungal VOC exposure also altered leaf VOC emissions, enriching alkanes, esters and monoterpenes. Initially, leaf VOC emissions were fungal lifestyle-specific but they converged over time, indicating systemic signal integration of belowground signals. These findings demonstrate that trees can discriminate “friend-versus-foe” through VOCs alone, extending pattern-recognition theory beyond contact-dependent cues. Multivariate analyses suggested organ-specific chemical strategies: roots function as chemosensors decoding fungal volatilomes, while systemic adjustments shape aboveground VOC profiles. Understanding the plant response to fungal VOCs may offer potential for developing early pathogen diagnostics and further elucidate the volatile-mediated plant-fungal interactions.

## Introduction

Plant-fungal interactions are an essential component of the structure and processes of terrestrial ecosystems (Philippot et al. 2013). Despite their microscopic nature, fungi play a crucial role in characterizing plant physiology, nutrient cycling, and robustness of the ecosystems (Lehmann and Kleber 2015; Heijden and Hartmann 2016; Bradford et al. 2016). The interactions between plants and fungi generally include three types: mutualistic, pathogenic and saprotrophic, each of them is able to regulate plant growth and development in a unique manner (Priyashantha et al. 2023). Whereas mycorrhizal fungi help plants stimulate acquisition of nutrients and stress endurance (Augé 2001; Smith et al. 2010; van der Heijden et al. 2015), pathogens impair host performance by triggering immune responses or transmitting effectors that interfere with host physiology (Doehlemann et al. 2017; Shao et al. 2021), and saprophytes help by decomposing and reforming the soil structure (van der Wal et al. 2013; Baldrian 2017; Clocchiatti et al. 2020). Plant roots are colonized by diverse fungal communities that vary significantly across environmental contexts, including soil properties, geographic factors, and biotic interactions (Jiang et al. 2020; Lin et al. 2024). Controlled experiments demonstrate that specific environmental factors, such as neighboring biotic elements and abiotic conditions, directly influence fungal assemblages associated with plant roots (Mony et al. 2021; Mészárošová et al. 2024). This established neighboring effect raises a deeper, mechanistic question: how do plants differentiate between their fungal neighbors, particularly before direct contact is made?

Volatile organic compounds (VOCs), a class of small airborne metabolites, represent a possible mechanism for pre-contact recognition in fungal-plant interactions. VOCs can diffuse through both soil airspaces and aqueous phases, with apolar VOCs being transmitted more rapidly in liquid environments due to their lack of hydration spheres (Weisskopf, Schulz and Garbeva 2021), thereby carrying chemically rich information over distances (Peñuelas et al. 2014). These compounds are important messengers in cross-kingdom communication among plants, fungi, and other microorganisms to convey information such as identity, physiological state, or potential dangers (Pichersky and Gershenzon 2002; Escobar-Bravo et al. 2023). Fungal VOC profiles are highly diverse and lifestyle-specific (Guo et al. 2021), and different effects have been observed: some species emit volatiles that promote plant growth or defense priming (Sharifi and Ryu 2021), but others also release suppressive or inhibitory signals (Bitas et al. 2013). Sesquiterpenes (SQTs), a subclass of terpenoids within the broader VOC family, are particularly prominent in fungal emissions and plant responses. These C15 compounds often mediate lifestyle-specific interactions, such as promoting root growth in mutualistic ectomycorrhizal associations via apolar SQTs produced by fungi like *Laccaria bicolor* (Ditengou et al. 2015) or acting as antimicrobial agents in pathogenic defenses (Sánchez-Fernández et al. 2016). These results suggest that plants may derive ecological background information from volatile cues and respond accordingly to improve survival and ecological flexibility. By incorporating SQTs into volatile bouquets, plants and fungi can fine-tune pre-contact recognition, potentially enhancing ecological adaptability in dynamic soil environments.

While VOC-mediated pre-contact recognition offers clear ecological advantages, plants still retain a robust contact-dependent recognition system. Pattern recognition receptors (PRRs), such as LysM-type receptors, detect fungal-associated molecular patterns like chitin and activate immune responses through MAPK signaling cascades (Spoel and Dong 2012; Liu et al. 2023). Recent studies show these receptors also fine-tune responses to modulate between defense and symbiosis, depending on context and dosage (Tan et al. 2025). VOC-mediated signaling may offer a complementary early-warning layer to this classical MAMP-based recognition. Additionally, under fungal challenge, plants often upregulate and emit theirselves sesquiterpenes that can function both as antimicrobial agents and intra-plant signals (Taniguchi et al. 2014; Frank et al. 2021; Laupheimer et al. 2023). However, the chemical overlap between fungal-derived and plant-derived sesquiterpenes creates a challenge: how can plants differentiate fungal signals from their own endogenous volatile production when molecules are structurally similar?

To investigate the capacity of plants to resolve such ambiguity and assign ecological value to fungal VOCs, we selected *Populus × canescens* as a plant model. This species is known for forming ectomycorrhizal symbioses with *Laccaria bicolor*, a well-characterized mutualist that influences root development and suppresses jasmonate signaling via the MiSSP7 effector (Felten et al. 2009; Daguerre et al. 2020). As representatives of other lifestyles, we included the saprotroph *Postia placenta*, involved in lignocellulose decay (Martinez et al. 2009), and the necrotrophic pathogen *Heterobasidion annosum*, known for aggressive root and stem infection (Hu et al. 2020; Tudoran et al. 2024). These fungi form ecologically distinct relationships with poplar and are known to produce lifestyle-informative VOCs (Müller et al. 2013; Guo et al. 2021).

Despite accumulating evidence of VOC-mediated plant-fungal communication, such as fungal volatiles reprogramming root architecture in ectomycorrhizal systems (Ditengou et al. 2015) and inducing short-term growth responses (Jiang et al. 2021), several questions remain unanswered. First, can plants reliably respond differentially to fungi with fundamentally different lifestyles (pathogenic, mutualistic, saprotrophic) using belowground volatile cues alone under standardized experimental conditions, particularly when distinguishing complex bouquets rather than individual compounds (Weisskopf, Schulz and Garbeva 2021)? Second, do these lifestyle-specific signals trigger distinct and predictable VOC responses in aboveground plant parts, and how are they spatially coordinated between roots and leaves (organ-specific vs. unified strategies)? Third, how do VOC-triggered responses unfold temporally - are they transient or sustained, and do they evolve from initial perception to long-term systemic adjustments (Bruisson et al. 2023)?

To address these questions, we employed a non-contact ‘pot-in-pot’ system that excluded direct physical interaction between plant roots and fungi and focused exclusively on volatile signaling. *P. × canescens* roots were exposed to VOCs from each fungus, and both root-zone and leaf emissions were tracked over time using GC–MS analysis. Multivariate statistical was employed to assess treatment discrimination and identify key discriminatory VOCs.

We tested three hypotheses: (I) poplar responses to fungal VOCs are dependent on fungal lifestyle, resulting in distinct volatile compositions in belowground and aboveground compartments; (II) These chemical interactions will exhibit temporal evolution from initial discrimination to adaptive integration over time, consistent with organ-specific and time-dependent effects observed in fungal volatile studies (Moisan et al. 2020; Singh et al. 2021; Olimi et al. 2025); and (III) sesquiterpenes dominate the discriminatory volatile classes in these interactions. By demonstrating that trees interpret airborne fungal signals before hyphal contact, this work widens the concept of pattern recognition beyond physical contact, offers a basis for non-invasive VOC-based diagnostics for early pathogen detection, and contributes to our understanding of plant-fungal chemical communication with implications for forest health monitoring.

## Materials and Methods

### Plant and microbial materials and growth conditions

This experiment used wild-type *Populus × canescens* (INRA clone 717–1B4). For propagation, plantlets from the established sterile stock were sectioned into segments under a laminar flow hood. These segments were then cultured in glass jars containing half-strength Murashige & Skoog (MS) medium, which had been adjusted to a pH of 5.8, and contained 30 g L^-1^ sucrose and 7 g L^-1^ agar (Duchefa Biochemie, Haarlem, Netherlands). The glass jars containing the plants were placed in a growth chamber at 20/16 °C (day/night), with 10-hour photoperiod (100 µmol m-² s^-1^ PPFD), for two months. Following the *in vitro* stage, the rooted plantlets were carefully transferred to pots containing an autoclaved substrate mixture (soil:vermiculite:perlite at a ratio of 3:1:1, v:v:v) and then all the plants were moved to a sun simulator (Kaling et al. 2014) with the following conditions: day/night temperature of 22/16 °C, relative humidity of 60/80% and light conditions of 10/14 h, 150 µmol m⁻² s⁻¹ PPFD) for three months to acclimatize to the different chamber conditions. The plants in the tray were first half-covered with plastic covers to maintain high humidity; the covers were removed gradually within two weeks.

Three fungal strains were used in the experiments based on their ecological lifestyles: the pathogen *Heterobasidion annosum* (stock collection Karin Pritsch, EUS, Helmholtz Munich); the mycorrhizal fungus *Laccaria bicolor* (S238N), and the saprotrophic *Postia placenta* (stock collection Karin Pritsch, Helmholtz Munich, Germany). The fungal cultures were grown in 35 mm × 10 mm culture dishes (Corning 430588, Corning Inc., NY, USA), which contained a modified Melin-Norkrans (MMN) medium (1% glucose, 0.25% NH₄-tartrate, 0.025% (NH₄)₂SO₄, 0.05% KH₂PO₄, 0.015% MgSO₄·7H₂O, 0.005% CaCl₂·2H₂O, 0.0025% NaCl, 0.01% FeCl₃, 0.001% thiamine hydrochloride and 1% Gelrite at a pH of 5.2 (Guo et al. 2021).

### Pot-in-pot exposure system and experimental setup

A custom ‘pot-in-pot’ (PiP) system (Fig. 1a) was developed to expose poplar roots to fungal VOCs without direct contact. The lower 2 cm of the upper pot was drilled with 28 evenly distributed 1 mm diameter holes to allow gas exchange between the upper and lower pots. Each PiP system consists of an upper pot containing 180 g of substrate (soil: vermiculite: perlite mixture at a ratio of 3:1:1, by volume) and a lower pot for the medium and fungi. All components were wrapped in aluminum foil and autoclaved at 121 °C for 20 min, two days prior to transferring the plants from the glass jar to the PIP system.

**Figure 1.**
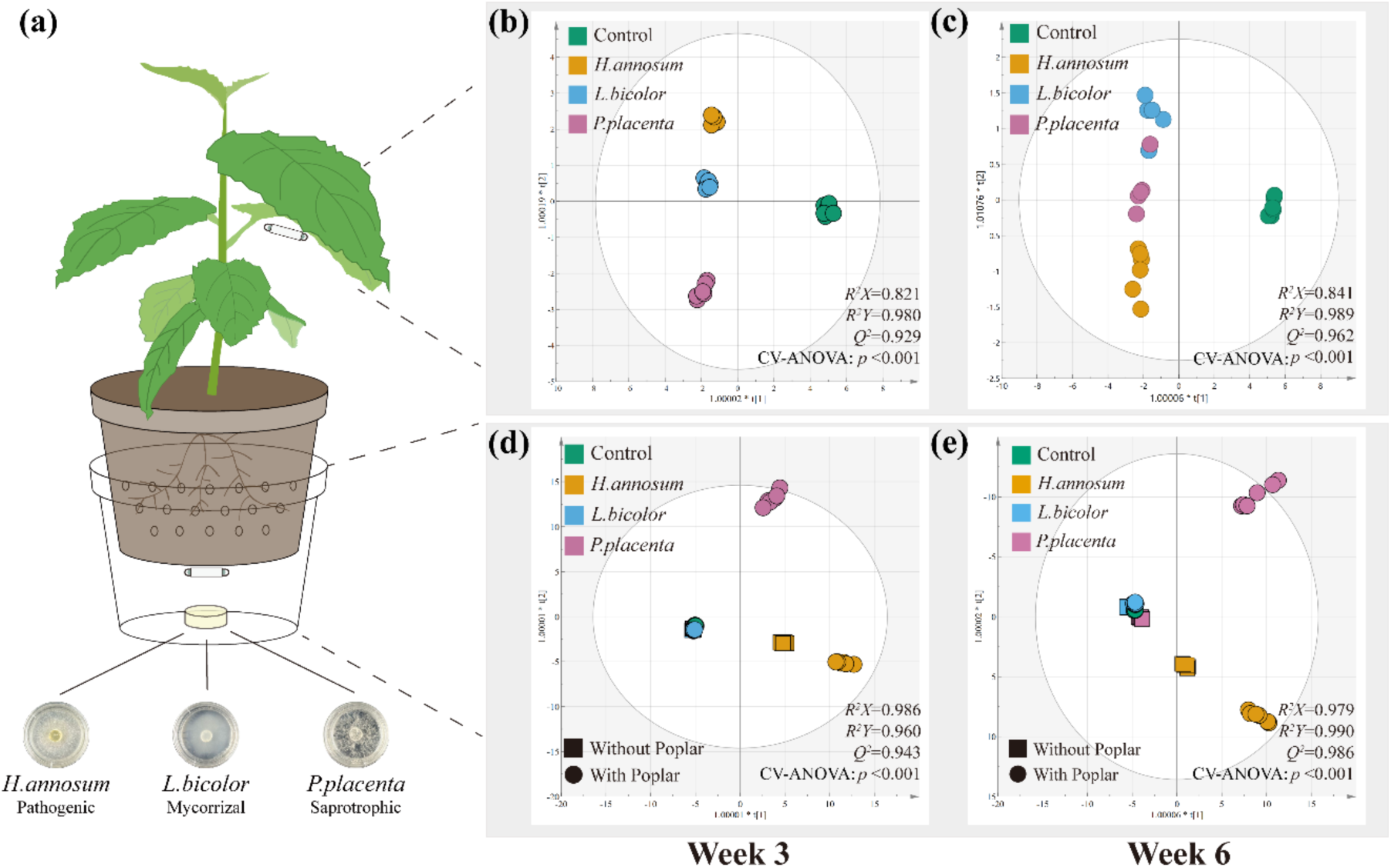
Experimental design and analysis of volatile organic compound (VOC) profiles in poplar exposed to volatiles from functionally diverse fungi. (a) Schematic showing poplar roots (upper pot) and fungal culture plates (lower pot) sharing headspace through perforated (upper) pot (28 × 1 mm holes) without physical contact. Fungal lifestyle: *H. annosum* = pathogenic; *L. bicolor* = mycorrhizal; *P. placenta* = saprotrophic. (b, c) OPLS-DA score plots of aboveground (leaf) VOC emissions at weeks 3 (b; *R²X* = 0.821, *R²Y* = 0.980, *Q²* = 0.929) and 6 (c; *R²X* = 0.841, *R²Y* = 0.989, *Q²* = 0.962) of co-cultivation. (d, e) OPLS-DA score plots of belowground VOC emissions at weeks 3 (d; *R²X* = 0.986, *R²Y* = 0.960, *Q²* = 0.943) and 6 (e; *R²X* = 0.979, *R²Y* = 0.990, *Q²* = 0.986); round symbols = with poplar, square symbols = without poplar. Each point represents one biological replicate (n = 6 per treatment). Model components were: aboveground VOCs - 3 predictive components (PC) (week 3, panel b) and 3 PC (week 6, panel c); belowground VOCs - 6 PC (weeks 3 and 6, panels d and e). Model parameters indicate good fit and predictive ability, with all models validated by permutation testing and cross-validation. All models were significant (*p* < 0.05, CV-ANOVA).

The experiment followed a 2 × 4 factorial design. The first factor was the presence or absence of a grey poplar plant in the upper pot. The second factor consisted of 3 fungal treatments and 1 control in the lower pot: a fungus-free MMN medium control (blank), and treatments containing fungi: *H. annosum*, *L. bicolor* and *P. placenta*. Each treatment had six biological replicates (n = 6), making a total of 48 PIP systems. The replicates of each treatment group were placed together in a single tray. To minimize position effects, different trays with poplar were systematically rotated within the climate chamber clockwise every three days.

To ensure continuous and active VOC emissions, the fungal plates in the lower pots were replaced weekly or when contamination on the fungal plate was observed. The fresh plates were pre-cultivated for different periods of time (*P. placenta*: 4 days; *H. annosum*: 8 days; *L. bicolor*: 28 days) to allow the mycelium to fully colonize the medium surface prior to replacement.

### VOC collection

Volatile organic compounds (VOCs) were collected at two time points: three weeks and six weeks after the onset of exposure of roots to fungal VOC emissions. For the belowground collection of VOCs from the root-zone headspace (which includes emissions from both plant roots and fungal cultures), a polydimethylsiloxane (PDMS) Twister (Gerstel GmbH, Mülheim an der Ruhr, Germany) was magnetically attached to the exterior bottom surface of the upper pot, directly above the fungal culture, and positioned in the headspace of the lower pot. The pot junction part was sealed to create a gas-tight seal, and volatiles were collected for 16 hours. This sampling approach captured the combined chemical environment, including fungal emissions and any root-derived compounds, allowing characterization of the complete root-zone VOC profile that plants experience during fungal exposure. Fungal monoculture treatments served as controls to distinguish fungal-derived compounds from plant-specific responses.

Aboveground VOC collection was performed by enclosing the entire aerial parts of each plant in a sealed 2 L Teflon bag, creating a gas-tight space physically separated from the root-zone, ensuring measured volatiles originated from aboveground plant tissues. A PDMS Twister was attached to the abaxial surface of a mature leaf. Volatiles were sampled for two hours under ambient light conditions to obtain the leaf VOC profile. After collecting, all Twisters were stored in sealed vials in a fridge at 4 °C until analysis.

### VOC analyses

Volatile organic compounds were analyzed on a GERSTEL Thermal Desorption Unit (TDU) with a Cooled Injection System (CIS 4), coupled to an Agilent 7890A gas chromatograph (GC) and a 5975C mass selective detector (MSD). Prior to analysis, an internal standard (860 pmol of δ-2-carene in hexane) was spiked onto each Twister. Each Twister was then thermally desorbed in the TDU from 30 °C (0.1 min) to 270 °C (2 min) at a rate of 280 °C/min. Desorbed analytes were cryofocused in the CIS at -50 °C before being injected in splitless mode. For injection, the CIS was heated to 270 °C (2 min) at 12 °C s^-1^. The transfer line was held at 250 °C.

Gas chromatographic separation was performed on a DB-5MS column (70 m × 0.25 mm i.d. × 0.25 µm film thickness; Agilent J&W) with helium as the carrier gas at a constant flow of 1.0 mL·min⁻¹. The GC oven program was as follows: 40 °C, ramped at 10 °C·min⁻¹ to 130 °C (held 5 min), then at 80 °C·min⁻¹ to 175 °C, then at 2 °C·min⁻¹ to 200 °C, then at 4 °C·min⁻¹ to 220 °C, and finally at 100 °C·min⁻¹ to 300 °C (held 5 min), for a total run time of 37.86 minutes.

The MSD was operated in electron-impact (EI) mode at 70 eV, with source and quadrupole temperatures at 230 °C and 150 °C, respectively. Data were acquired in a combined SIM/scan mode after an 8.3 min solvent delay. Full scan mode monitored m/z 35–250. Two SIM groups were used to enhance sensitivity for monoterpenes (m/z 93, 121, 136; starting at 8.3 min) and sesquiterpenes (m/z 93, 161, 204; starting at 14.9 min). Isoprene could not be measured due to its extremely high volatility and insufficient retention on the PDMS Twister sorbent material.

Compounds were identified by comparing their mass spectra and retention indices with the NIST 2020 spectral library and authenticated standards. Quantification was performed using external calibration curves. Emission rates were calculated based on the peak area of each compound relative to the peak area of the internal standard (δ-2-carene) and further normalized according to sampling duration and either aboveground leaf area or belowground Petri dish area (Ghirardo et al. 2011, 2020).

### Statistical Analysis

All statistical analyses were conducted on the time-resolved volatilomics data, which were processed as two separate matrices: the belowground dataset (56 compounds, Supplemental Table S1) and the aboveground dataset (57 compounds, Supplemental Table S2). Prior to analysis, all emission rate data were log₁₀-transformed to approximate a normal distribution, then mean-centered and Pareto-scaled. An exploratory principal component analysis (PCA) was initially performed on each dataset to inspect data variation and identify potential outliers. One sample falling outside the 95% Hotelling’s T² confidence ellipse was excluded from further analysis.

For both the aboveground and belowground VOC datasets, separate Orthogonal Partial Least Squares Discriminant Analysis (OPLS-DA) were then performed using SIMCA-P (v. 13.0.3.0, Umetrics, Umeå, Sweden). In these supervised models, the respective matrix of VOC emission rates served as the independent variables (X), while the categorical variable indicating the fungal treatment (*Heterobasidion annosum*, *Laccaria bicolor*, *Postia placenta* or control) was set as the dependent variable (Y). The performance and reliability of the OPLS-DA models were assessed using cumulative explained variance (*R²Y*), predictive ability (*Q²Y*), root mean square error of estimation (RMSEE) and root mean square error of cross-validation (RMSECV). Model significance was tested using a cross-validated analysis of variance (CV-ANOVA, *P* < 0.05) (Eriksson, Trygg and Wold 2008; Eriksson et al. 2013). OPLS-DA score plots were then used to illustrate the differentiation of poplar VOC profiles based on fungal exposure.

Further statistical analyses and data visualizations were conducted in R (v.4.3.3; R Core Team, 2023). For normally distributed data, differences between groups were tested using a one-way ANOVA followed by a Tukey’s HSD post hoc test (*p* < 0.05). For univariate comparisons of non-normally distributed transformed data, a Kruskal–Wallis test was used, with Bonferroni-adjusted post hoc comparisons for emission-rate data. Hierarchical clustering analysis was performed using complete linkage with Euclidean distances. The resulting patterns of VOC abundance across treatments were visualized as heatmaps using the ‘pheatmap’ package in R. Non-metric multidimensional scaling (NMDS), an ordination method, was used to visualize VOC community structure based on Bray-Curtis dissimilarity using the ‘vegan’ package in R. The homogeneity of multivariate dispersions was tested using the ‘betadisper’ function before conducting PERMANOVA for significance testing, with Holm adjustment applied to pairwise comparisons. β-diversity was calculated as the mean Bray-Curtis dissimilarity between samples within each treatment group compared to all other treatment groups (between-group beta diversity), averaged across all pairwise sample comparisons and expressed as mean ± SEM.

## Results

### Plant-mediated detection of fungal lifestyle through distinct fungal VOC signatures

Poplar exposure to VOCs from contrasting fungal lifestyles showed distinct below- and above-ground volatile emission responses (Fig.1). To quantify separation, we applied OPLS-DA analysis of VOCs detected in the headspace between the air-permeable pot and the fungal culture (56 compounds; see Supplementary Table S1), and in the above-ground plant parts (57 compounds; see Supplementary Table S2) revealed that exposure of poplar roots to fungal VOCs induced significant (*p*<0.001, CV-ANOVA) distinct changes in the root-zone headspace composition compared to VOCs emitted by roots alone or by fungal mycelia in the absence of plant contact (Fig. 1). These VOC fingerprints clearly distinguish between pathogenic (*H. annosum*), mycorrhizal (*L. bicolor*) and saprophytic (*P. placenta*) interactions, as well as between the control groups (monocultures or root emissions without fungal contact), and between each other (Fig. 1b–e).

At week 3, the three lifestyle treatments formed distinct clusters with clear separation along the first predictive component (PC) (Fig. 1b). By week six, differences among fungal treatments decreased, though all co-cultivation treatments remained distinguishable from controls (Fig. 1c).

In contrast to the temporal changes observed aboveground, the VOC patterns in the headspace between the perforated pot (allowing air exchange) and the fungal culture remained temporally stable. At week three, the OPLS-DA showed distinct separation among fungal lifestyles and controls (Fig. 1d). Importantly, for each fungal species, the presence of poplar roots shifted VOC profiles away from their respective monoculture baselines, demonstrating that plant presence actively alters the chemical environment rather than simply reflecting inherent differences between fungal lifestyles. The pathogenic *H. annosum* exhibited the greatest difference between its monoculture and co-culture states, indicating the strongest plant-fungus chemical interaction. At week six, this plant-mediated separation became even more pronounced (Fig. 1e). The distance between monoculture and co-culture conditions increased or persisted over time for the fungi, with minimal change for *L. bicolor* (Fig. 1d, e). *H. annosum* co-cultures remained uniquely separated from all other treatments, with greater magnitude of shift compared to mycorrhizal or saprophytic interactions. High compound overlap between time points (*L. bicolor* 100%, *H. annosum* 93%, *P. placenta* 84%) confirmed temporal consistency in these interaction-specific, rather than simply fungal-specific, volatile profiles (Supplementary Fig. S1).

Together, the three fungal species representing different lifestyles and the responses from different root-zone and leaf responses showed different fingerprints of VOCs profile: the belowground VOC signatures remained temporally stable and treatment-specific, while the leaf VOC profiles gradually became more similar over time, but still maintained significant differences from the control group.

### Belowground fungal lifestyle determines plant-modulated volatile signatures with distinct chemical class specialization

To identify the specific compounds driving these discriminant patterns, we performed a hierarchical cluster analysis of the individual volatile compounds. Across all treatments and time points, we detected 56 VOCs in the root-zone headspace, mainly sesquiterpenes (84%). Hierarchical clustering identified specific compound clusters that distinguished the three fungal lifestyles in co-cultivation with poplar (Fig. 2). Compound-overlap analysis showed minimal sharing among lifestyles (typically <5 shared compounds): in monoculture at week three, *H. annosum* exhibited 15 unique compounds, whereas *L. bicolor* and *P. placenta* showed 2 and 0, respectively (Fig. S2a). Upon co-cultivation with poplar, all fungi emitted broader and more complex signatures (Fig. S2b); *H. annosum* maintained the highest uniqueness (18 compounds), and *P. placenta* increased from 0 to 14. These patterns persisted at week six, with *H. annosum* reaching 19 unique compounds and *P. placenta* maintaining 13 (Fig. S2c, d).

**Figure 2.**
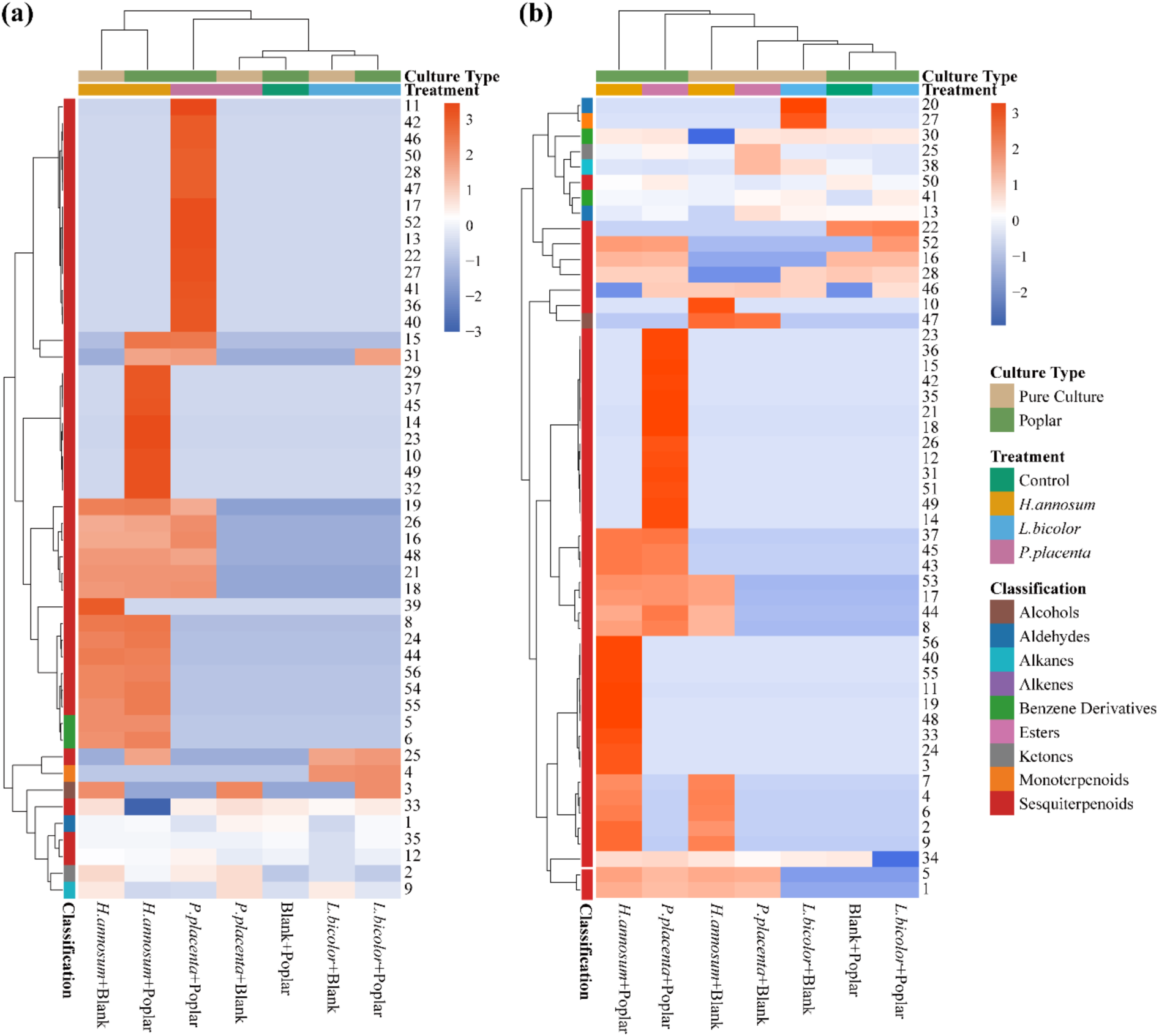
Hierarchical clustering reveals distinct belowground VOC profiles under contact-free fungal co-cultivation. (a, b) Bi-dimensional clustered heatmaps showing belowground Z-score normalized volatile compound abundances at weeks 3 (a) and 6 (b). Rows represent individual compounds (IDs in Supplementary Table 1); columns show treatment groups including pure fungal cultures (*Heterobasidion annosum*, *Laccaria bicolor*, *Postia placenta*) and co-cultures with poplar. Hierarchical clustering used Euclidean distance and complete linkage. Left color bar indicates chemical classification; top bars show culture type and treatments. Color intensity (blue to orange red) reflects Z-score values. Data represents mean values from n = 6 biological replicates per treatment.

Temporal comparisons indicated substantial consistency in VOC emissions: 30/56 compounds (54%) showed consistent responses across weeks three and six, while 16 increased and 10 decreased in abundance. Pairwise Spearman correlation matrices revealed denser, positively correlated modules under poplar exposure - most pronounced for *H. annosum* and *P. placenta* - consistent with coordinated metabolic responses rather than random additions (Figs. S3–S4).

Treatment-specific changes were sesquiterpene-dominated (over 80%). In co-cultivation, additional sesquiterpenes were detected with *H. annosum* (e.g., isoledene, cedrene), and *P. placenta* shifted from weak monoculture emissions to diverse sesquiterpenes (e.g., silphinene, β-caryophyllene). In contrast, exposure to *L. bicolor* showed limited changes in overall composition under co-cultivation. Control treatments (plants without fungal exposure and fungi without plants) exhibited low VOC emissions.

### Compartment-specific chemical strategies reveal temporal convergence in aboveground volatile responses

To test whether fungal volatiles detected from below-ground trigger systemic aboveground responses, we analyzed 57 volatiles released from *P. × canescens* leaf tissue. Hierarchical cluster analysis of volatiles revealed distinct, fungal-specific leaf emissions (Fig. 3), showing systemic chemical responses to different fungal VOCs exposed below-ground. Although VOCs detected in the belowground compartment were dominated by sesquiterpenes (84%), the aboveground emissions exhibited higher chemical diversity, with alkanes (35%) and monoterpenes (25%) being the most abundant compound classes (isoprene could not be measured due to methodological limitations). In contrast to the below-ground compartment, above-ground VOC profiles showed increasing similarity across treatments over time, as evidenced by NMDS (Fig. 5c, d).

**Figure 3.**
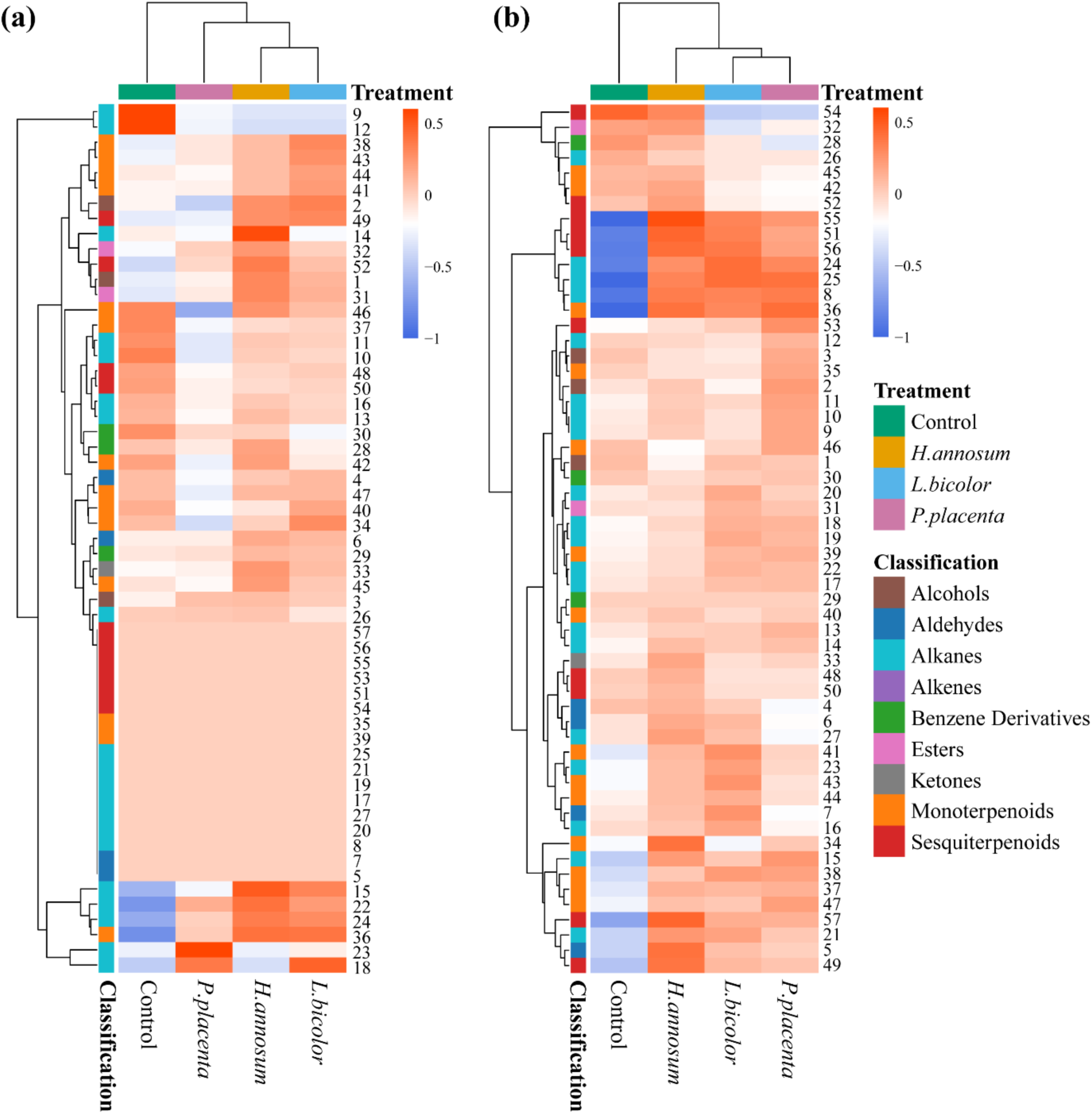
Hierarchical clustering of aboveground leaf VOC profiles in poplar exposed to fungal volatiles belowground. (a, b) Heatmaps showing Z-score normalized volatile compound abundances from poplar leaves at weeks 3 (a) and 6 (b) of co-cultivation. Rows represent individual compounds (IDs on right, see Supplementary Table 2); columns show treatments where plants were exposed to volatiles from different fungal cultures (*H. annosum*, *L. bicolor*, *P. placenta*) or control medium via the non-contact pot-in-pot system. Hierarchical clustering is performed using Euclidean distance and complete linkage. Left color bars indicate chemical classification of compounds; top color bars show fungal treatments. Color intensity (blue to orange) reflects Z-score values. Data represents mean values from n = 6 biological replicates per treatment.

At week three of exposure, the heatmap analysis revealed pronounced lifestyle-specific clustering patterns in aboveground volatile profiles (Fig. 3a). The pathogenic *H. annosum* induced distinctive systemic signature in the leaves, characterized by broad upregulation across chemical classes, particularly esters such as (Z)-3-hexen-1-ol acetate (compound 31), which increased to 6.83 compared to 4.20 nmol m⁻² s⁻¹ in controls. The ectomycorrhizal *L. bicolor* induced a selective leaf response, with elevated monoterpenoids but little changes in alkanes and aldehydes. In contrast, the exposure of saprotrophic *P. placenta* volatiles did not lead to a strong aboveground leaf response, as seen from the minimal deviation from control levels across most chemical classes with many compounds decreased in abundance compared to control.

By week six, fungal VOC-discrimination decreased, as evidenced by reduced dendrogram distances in the hierarchical clustering analysis (Fig. 3b). The treatment-specific branches from week three shifted toward more overlapping patterns, with reduced separation between groups. This convergence appeared particularly in the monoterpenoid and sesquiterpenoid classes, where initial distinct signatures transitioned to more consistent levels across treatments. However, some sesquiterpenes (compounds 49, 57) and monoterpenes (compound 34) showed higher abundance in the *H. annosum* treatment relative to others. For the overlapping compound bouquets, different lifestyle fungi still led to distinct patterns.

Despite this temporal convergence, all fungal treatments were clearly distinguishable from control conditions at both time points, but their differences increased at week six, as seen in the dendrogram distances for sesquiterpenes and alkanes, indicating a remarkable systemic impact of belowground volatile exposure on leaf metabolism. This temporal dynamic in aboveground responses contrasted markedly with the persistent lifestyle-specific discrimination observed in belowground volatile profiles throughout the experimental period.

### Emission rate quantification reveals compartmentalized VOC emissions with opposing temporal dynamics

Quantitative emission profiling revealed spatial specialization and temporal divergence in plant-fungal chemical interactions (Figs. 4,5). Root-zone headspace emissions spanned a range of 0.5-32.1 pmol m⁻² s⁻¹ and exhibited fungal lifestyle-dependent signatures. At week three, co-cultivation with *H. annosum* resulted in the highest belowground emissions (31.6 pmol m⁻² s⁻¹), consisting of 98% sesquiterpenoids (Fig. 4a). *Laccaria bicolor* showed low emissions both in monoculture and in co-cultivation with poplar (1.71 pmol m⁻² s⁻¹). *Postia placenta* increased emissions from nearly zero in monoculture to 9.80 pmol cm⁻² h⁻¹ when co-cultivated with poplar by week six (Fig. 4b).

**Figure 4.**
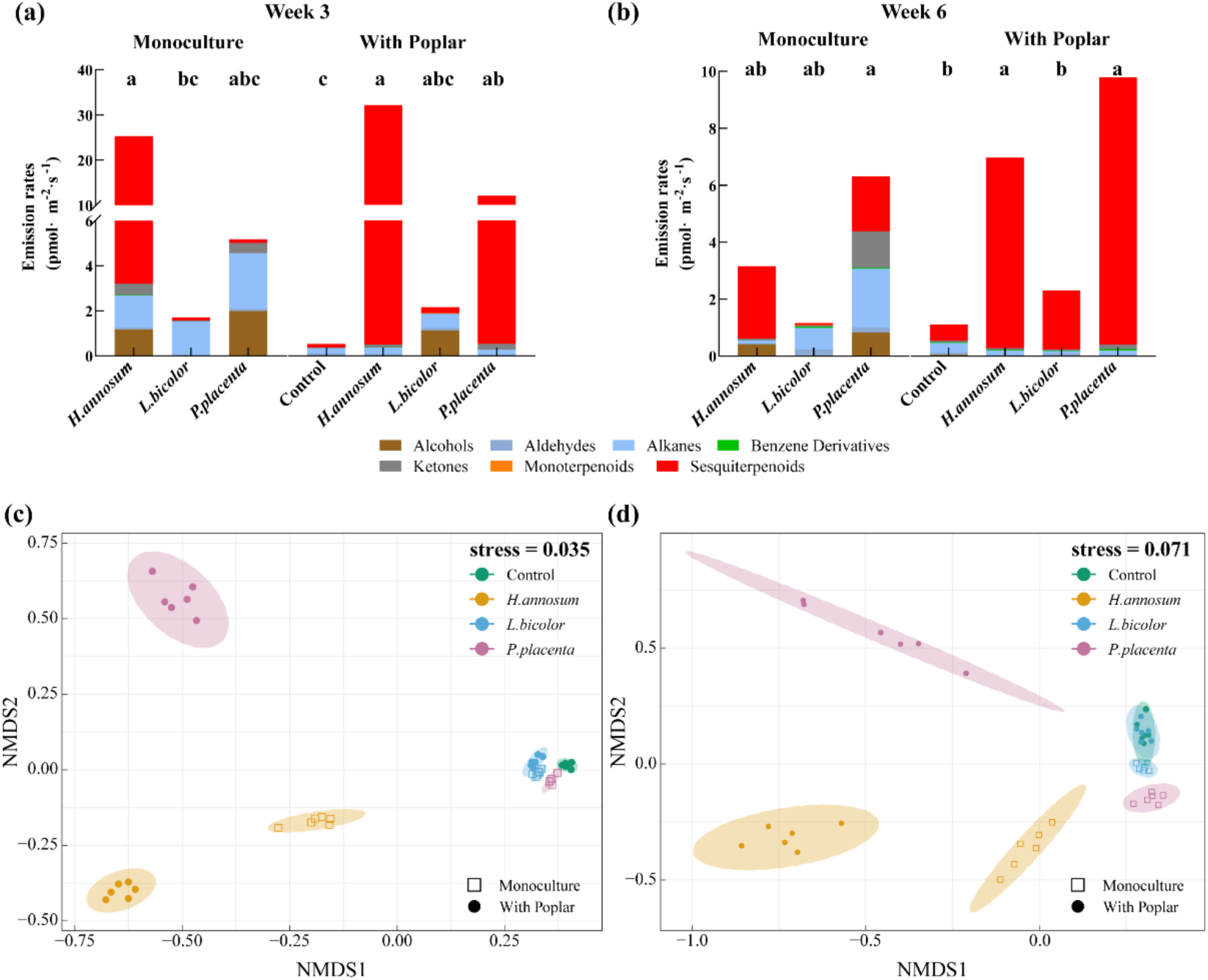
Effects of contact-free fungal co-cultivation on belowground VOC emission profiles and NMDS of VOC profiles over time. (a, b) Stacked bar plots depicting total VOC emission rates (pmol m⁻² s⁻¹) at week 3 (a) and week 6 (b), categorized by chemical functional groups. Left panels show emissions from fungal monocultures; Right panels illustrate VOC emissions from the root-zone headspace (including emissions from both poplar roots and fungal cultures) in a contact-free pot-in-pot system where poplar plants were exposed to fungal VOCs. Co-cultivations include *Heterobasidion annosum* (*H. annosum*), *Laccaria bicolor* (*L. bicolor*), and *Postia placenta* (*P. placenta*), with a monocultural controls. Different lowercase letters above denote significant differences among treatments (P < 0.05, Kruskal-Wallis test followed by Dunn’s post hoc test with Bonferroni correction). (c, d) non-metric multidimensional scaling (NMDS) ordination plots showing β-diversity of belowground VOC communities at week 3 (c) and week 6 (d), based on Bray-Curtis dissimilarity indices. Points represent individual replicates, colored and shaped by treatment group as indicated in the legends. Ellipses (dashed for monocultures, solid for poplar exposures) enclose group clusters. Stress values reflect ordination fit (week 3: 0.035; week 6: 0.071). Overall treatment effects were assessed by PERMANOVA: week 3 – *F*₆,₃₄ = 12.96, *R²* = 0.696, *P* = 0.001; week 6 – *F*₆,₃₅ = 5.02, *R²* = 0.463, *P* = 0.001. Data is from n = 6 biological replicates

NMDS of belowground VOC profiles showed lifestyle-specific structuring across time points (Fig. 4c, d; model fit (stress) = 0.035 and 0.071, respectively). Treatment groups formed distinct, non-overlapping clusters, with stable patterns from week three to week six as clusters maintained relative positions and separation distances. Multivariate analysis showed significant treatment separation (PERMANOVA: week 3: *F₆,₃₄*= 725.86, *R²* = 0.992, *P* = 0.001; week 6: *F₆,₃₅* = 409.58, *R²* = 0.986, *P*= 0.001). β-Diversity (calculated as Bray-Curtis dissimilarity between samples within each treatment group) analysis showed declining within-group variability over time, with fungal treatments converging to low Bray-Curtis dissimilarities by week six, while controls retained higher heterogeneity (Table S3).

Leaf volatile emissions ranged from 4.6–7.5 nmol·cm⁻²·h⁻¹ and were dominated by esters (46– 68%) and monoterpenoids (Fig. 5a, b). Among the esters, (*Z*)-3-hexen-1-ol acetate emissions were higher in the *H. annosum* treatment (6.83 nmol·cm⁻²·h⁻^1^) compared to controls (4.20 nmol·cm⁻²·h⁻^1^) and other fungal lifestyles in week 3 (Fig. 5a), but methyl salicylate (MeSA) emission in control (2.29 nmol·cm⁻²·h⁻^1^) is higher than *H.annosum* (1.68 nmol·cm⁻²·h⁻^1^). By week 6, these emissions converged across treatments, with (Z)-3-hexen-1-ol acetate levels around 1 nmol·cm⁻²·h⁻¹ in all groups, while MeSA increased overall, with elevations particularly evident in pathogenic treatments (6.13 nmol·cm⁻²·h⁻¹) relative to ectomycorrhizal and saprotrophic groups.

**Figure 5.**
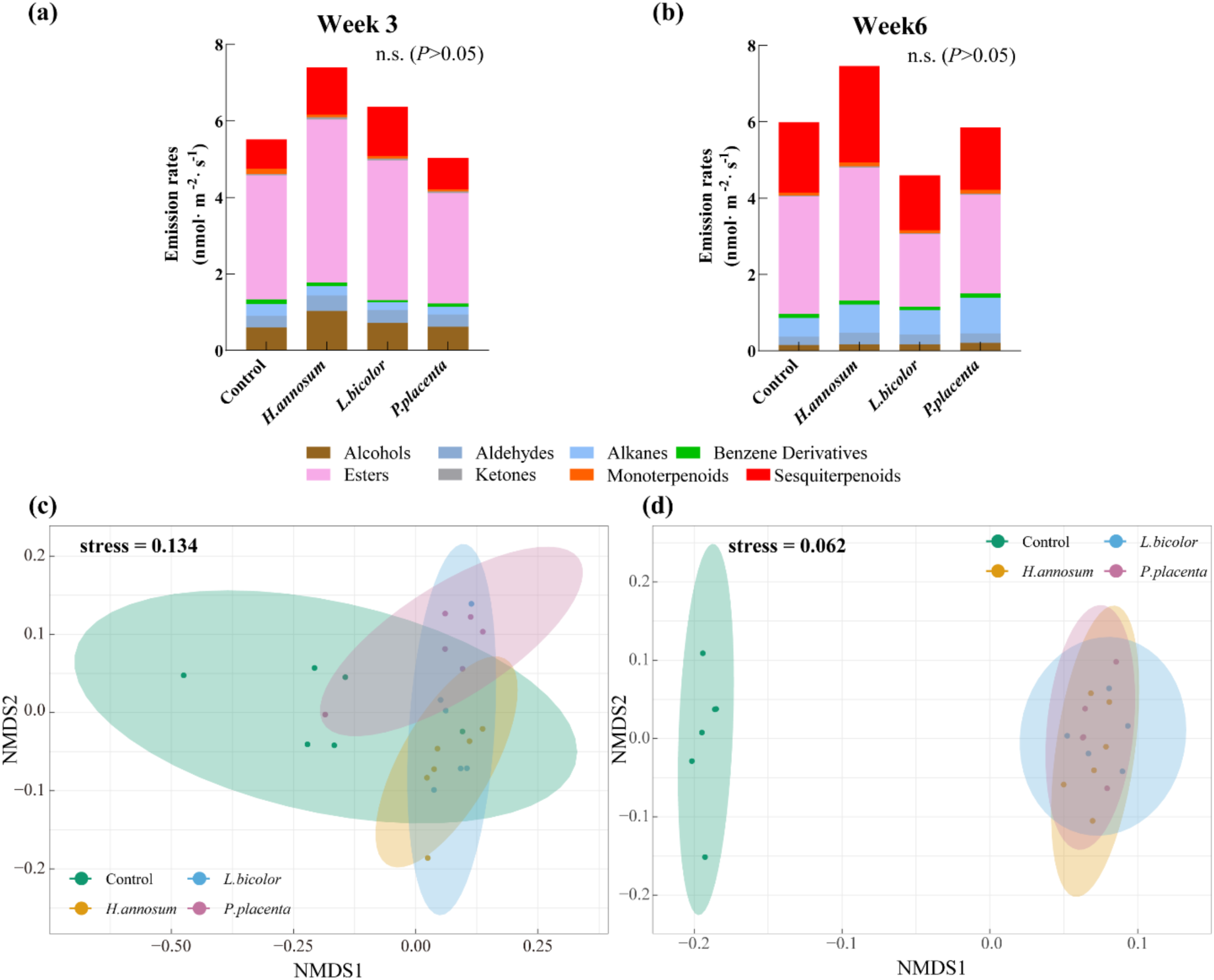
Effects of fungal VOC exposure on aboveground leaf VOC emission profiles and NMDS of VOC profiles over time. (a, b) Stacked bar plots depicting VOC emission rates at week 3 (a) and week 6 (b) of co-cultivation, categorized by chemical functional groups. Treatments include poplar plants without fungal exposure (Control) and those exposed to fungal VOCs from *Heterobasidion annosum* (*H. annosum*), *Laccaria bicolor* (*L. bicolor*), or *Postia placenta* (*P. placenta*) in a contact-free pot-in-pot system. No significant differences in emission rates were observed among treatments at either week 3 or week 6 (n.s., *P* > 0.05, Kruskal-Wallis test followed by Dunn’s post hoc test with Bonferroni correction). (c, d) Non-metric multidimensional scaling (NMDS) ordination plots showing β-diversity of aboveground VOC communities at week 3 (c) and week 6 (d), based on Bray-Curtis dissimilarity indices. Points represent individual replicates, colored and shaped by treatment group as indicated in the legends. Ellipses enclose group clusters. Stress values reflect ordination fit (week 3: 0.134; week 6: 0.062). Overall treatment effects were assessed by PERMANOVA: week 3 – *F₃,₂₀* = 3.51, *R²* = 0.345, *P* = 0.002; week 6 – *F₃,₁₈* = 6.00, *R²* = 0.500, *P* = 0.001. Data are means from n = 6 biological replicates

No significant differences in total leaf emission rates were observed among treatments at either time point. NMDS ordination showed temporal convergence (Fig. 5c, d; stress = 0.134 and 0.062, respectively). At week 3, treatments formed partially overlapping but distinguishable clusters (Fig. 5c). By week 6, groups exhibited substantial overlaps, with subtle separation from controls (Fig. 5d). Treatment effects were significant across time points (PERMANOVA: week 3: *F₃,₂₀* = 3.51, *R²* = 0.345, *P* = 0.002; week 6: *F₃,₁₈* = 6.00, *R²* = 0.500, *P* = 0.001). Fungal treatments remained distinguishable from controls (root: *P* < 0.001; leaf: *P* < 0.01) without physical contact.

## Discussion

Our study demonstrates that poplars can distinguish among a pathogenic, mutualistic, and saprotrophic fungi solely through sensing fungal VOCs, triggering a systematic response that alters aboveground leaf VOC emissions. This finding suggests a key mechanism by which plants navigate the complex chemical dialogue of the rhizosphere, where interactions are shaped not only by soluble root exudates that mediate mutualisms (Yu et al. 2022), but also by airborne signals that enable pre-contact recognition between fungi and plants (Ding et al. 2020; Feng et al. 2025). This finding aligns with the broader view that microbial volatiles function as inter-kingdom infochemicals (Schulz-Bohm, Martín-Sánchez and Garbeva 2017; Weisskopf, Schulz and Garbeva 2021)).

We addressed the question of whether plant responses to fungal VOCs are generalized or specific. Systemic leaf volatile emissions reflected lifestyle-specific, rather than generalized responses to fungal presence. The results showed strategy-resolved patterns rather than generalized stress responses. OPLS-DA and hierarchical clustering of leaf volatiles revealed distinct chemical signatures for each fungal lifestyle (Fig. 3). Pathogenic *H. annosum* induced a broad upregulation across multiple compound classes, ectomycorrhizal *L. bicolor* triggered selective monoterpenoid responses, and saprotrophic *P. placenta* did not cause remarkable changes compared to controls. These differentiated systemic patterns indicate non-additive outcomes and suggest specific plant recognition mechanisms at the plant–fungus interface, phenomena that are increasingly recognized in complex plant-microbe interactions where plants can produce synergistic benefits (Hestrin et al. 2019; Rawat et al. 2025) through deep molecular regulation (Gao et al. 2023).

Regarding temporal stability, belowground VOC profiles-maintained discrimination over six weeks, with consistent separation in NMDS ordinations (stress 0.035–0.071; Fig. 4c, d) and declining Bray-Curtis dissimilarity (PERMANOVA *P* = 0.001; Table S3). This persistence aligns with the belowground aspect of our hypothesis I, contrasting transient fluctuations observed in other systems, such as rapid protein phosphorylation changes in response to green leaf volatiles (Tanarsuwongkul et al. 2024) or short-term microbial gene expression shifts induced by fungal VOCs (Chang et al. 2024), and emphasizing sustained ecological strategies in our non-contact setup. On spatial coordination versus compartmentalization, responses were clearly divided: from root-zone sesquiterpene-dominated profiles (0.5–32.1 pmol m⁻² s⁻¹) were detected, while leaves shifted to alkane/ester/monoterpene blends (4.6–7.5 nmol m⁻² s⁻¹; Figs. 4a, b and 5a, b). This compartmentalization supports a model of localized sensing in roots and systemic integration in leaves, which aligns with the introduction’s emphasis on whole-plant coordination. This finding also corresponds with the aboveground temporal convergence predicted in hypothesis II, as evidenced by NMDS clustering (stress 0.134–0.062; Fig. 5c, d). Notably, the dominance of sesquiterpenes in roots (comprising 84% of compounds) partially meets hypothesis III, though leaf diversity suggests broader chemical classes contribute to systemic responses.

Our study revealed that exposure to the fungal pathogen *H. annosum* induced high belowground sesquiterpene emissions (31.6 pmol m⁻² s⁻¹ at week three, persisting through week six; Fig. 4a, b), including additional compounds such as isoledene and cedrene (Fig. 2a, b). This response may be mediated by rapid intracellular calcium (Ca²⁺) transients, a known early signaling event in plant-pathogen interactions that activates downstream defense pathways (Zhang, Du and Poovaiah 2014); however, direct quantification of these events was not performed. The sustained high sesquiterpene emissions is consistent with a barrier program that can be initiated in advance (Aratani et al. 2023). From a cost-benefit perspective, such chronic elevation represents an investment in preemptive barriers, reflecting a fundamental growth-defense tradeoff shaped by evolutionary optimization (Monson et al. 2022). This hypothesis is further supported by findings that species tend to maximize only one of several possible defense strategies (Sancho-Knapik et al. 2023). From an ecological perspective, this strategy of deploying high levels of constitutive antifungal terpenes has been shown to be a proven mechanism for preventing necrotrophic invasion, thereby contributing to forest pathogen resistance (Wang et al. 2024).

In contrast, the ectomycorrhizal fungus *L. bicolor* elicited consistently low emissions (1.71 pmol m⁻² s⁻¹; Fig. 4a, b). across all time points and showed minimal compositional changes upon co-cultivation. This restrained VOC profile does not suggest a lack of communication but rather reflects a highly efficient ’less is more’ signaling strategy. This is supported by studies showing that even low-abundance sesquiterpenes from *Laccaria* are potent enough to reprogram host root architecture (Ditengou et al. 2015). This targeted approach, possibly modulated by effectors like MiSSP7 (Daguerre et al. 2020), may optimize mutualistic establishment by avoiding the activation of broad-spectrum host defenses while maintaining effective chemical communication.

The saprotrophic *P. placenta* exhibited dramatic plasticity in response to plant presence, with VOC emissions increasing from near-zero monoculture levels to 11-12 pmol m⁻² s⁻¹ when co-cultivated with poplar, maintaining elevated emissions (∼9.80 pmol m⁻² s⁻¹) through week six (Fig. 4a, b). This response was accompanied by compositional shifts toward sesquiterpenes such as silphinene (Fig. 2a, b). Rather than indicating pathogenic behavior, this dynamic response likely reflects the opportunistic resource-sensing capacity typical of saprotrophic fungi, where metabolic investment scales with substrate availability (Marañón-Jiménez et al. 2021). The temporal increase suggests adaptive metabolic flexibility rather than aggressive host exploitation, consistent with the decomposer lifestyle that relies on detecting and efficiently utilizing available organic matter (Metz et al. 2025).

Exposure to fungal VOCs belowground induced systemic changes in leaf emissions aboveground. Initially, plants discriminated between fungal lifestyle, for instance ester upregulation under *H. annosum* (e.g., (*Z*)-3-hexen-1-ol acetate) (Fig. 3a, b). Over time, these initially distinct VOC profiles showed convergence patterns (PERMANOVA *P* = 0.001–0.002; Fig. 5c, d), reflecting a shift from acute GLV-type responses toward more persistent salicylate-linked signaling, as indicated by increased MeSA levels at week six. This temporal progression from immediate GLV responses to sustained MeSA signaling is consistent with plant defense signaling patterns, where early volatile responses transition to more persistent systemic signals over prolonged exposure (Frank et al. 2021; Laupheimer et al. 2023; Gong et al. 2023).

The present findings reveal volatile-mediated surveillance as a complementary early layer of plant-microbe interactions, extending beyond contact-dependent PRR-MAMP recognition (Medina-Puche et al. 2020). Perception may involve membrane-associated candidates and KAI2-related pathways proposed for volatile/karrikin sensing in other contexts(Bythell-Douglas et al. 2017); we did not assay receptors here, so genetic tests in *Populus* will be required in the future (Stirling et al. 2024). The shared terpenoids may complicate cue origin, yet enable pattern-level decoding of mixture features (Zeng et al. 2022). In fact, this ambiguity may be ecologically beneficial, enabling shared chemical channels for compatibility signaling and threat recognition (Kanchiswamy, Malnoy and Maffei 2015; Sharma, Anand and Kapoor 2017). Ecologically, such a system can contribute to microbiome assembly by allowing root VOCs to filter communities and facilitate feedbacks like neighbor facilitation (Gfeller et al. 2019; Mészárošová et al. 2024), though further studies are needed to validate these adaptive benefits.

Several limitations should be acknowledged. First, the simplified ‘pot-in-pot’ system, cannot replicate the spatial heterogeneity, real-soil gradients, and multi-species dynamics that influence VOC diffusion in natural environments (Parlin et al. 2022). Second, our six-week study window captures only early interaction dynamics but may overlook long-term acclimation processes (Beckman, Dybzinski and Tilman 2023). Third, the origin of individual compounds remain ambiguous without source tracing; future studies should employ stable isotopes to assign VOC to plant or fungal sources (Ghirardo et al. 2011; Honeker et al. 2023). Fourth, the bidirectionality of VOC signaling needs validation via reciprocal assays and genetic approaches (e.g., terpene synthase (TPS) knockouts). Additional methodological concerns include culture plate replacement or batch effects, VOC adsorption to experimental material, and differences in diffusion coefficients between sealed PIP systems and natural soil pore structures. To address these limitations future studies should incorporate mesocosms with diverse fungi, proteomics analysis to identify plant receptors, and field studies to evaluate the generality of these finding beyond *Populus*.

In conclusion, our study identified four key findings of plant volatile-mediated recognition: (1) Plants detect and discriminate among fungal lifestyles based solely on airborne VOC cues, demonstrating specific rather than generalized responses that likely confer evolutionary advantage by enabling early threat detection in complex rhizosphere environments. (2) Chemical signaling exhibited organ-specific patterns, with belowground responses maintaining discriminatory specificity while aboveground responses showed convergent patterns, indicating a balance between selective recognition and metabolic efficiency that extends beyond binary classification systems. (3) Functional compartmentalization was observed, with sesquiterpenes dominating the discriminatory volatile classes, where root systems served as primary sensory organs and leaf tissues as integrative response centers. These results establish volatile-mediated surveillance as a complementary early-stage recognition mechanism in trees, widening the concept of pattern recognition beyond physical contact and offering potential for VOC-based diagnostics in forest health monitoring.

## Acknowledgements

The authors thank Karin Pritsch (Helmholtz Munich, Research Unit Environmental Simulation) for provision of fungal strains and valuable advice and help with maintaining the fungal cultures.

## Author’s contributions

P.Z., M.R., and J.-P.S. designed research; P.Z. performed research with B.W. (GC-MS); P.Z. analyzed data; P.Z., M.R., A.G., and J.-P.S. interpreted data; and P.Z. wrote the paper with input from all authors

## Data Availability Statement

The data that support the findings of this study are available on request from the corresponding author. The data is not publicly available due to privacy or ethical restrictions.

## Supporting Information

### Supplemental Figures and Tables

**Supplementary Figure 1.**
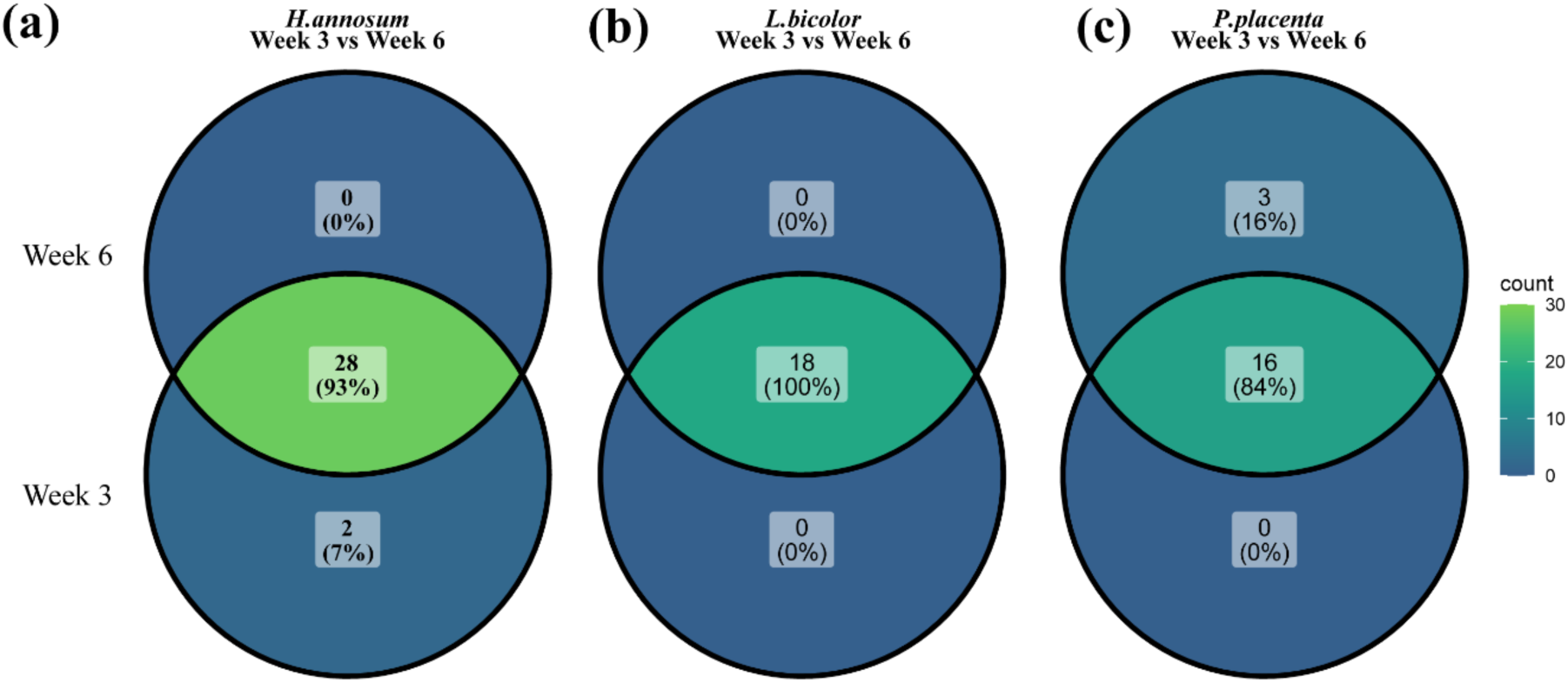
Venn diagrams showing temporal stability of VOC profiles in axenic fungal monocultures between Week 3 and Week 6. Monocultures of (a) *Heterobasidion annosum*, (b) *Laccaria bicolor*, and (c) *Postia placenta* were maintained axenically, without *Populus × canescens*. Numbers indicate the count of detected VOCs, with percentages showing the proportion of shared compounds between timepoints. Colors represent compound abundance according to the scale bar (right).

**Supplementary Figure 2.**
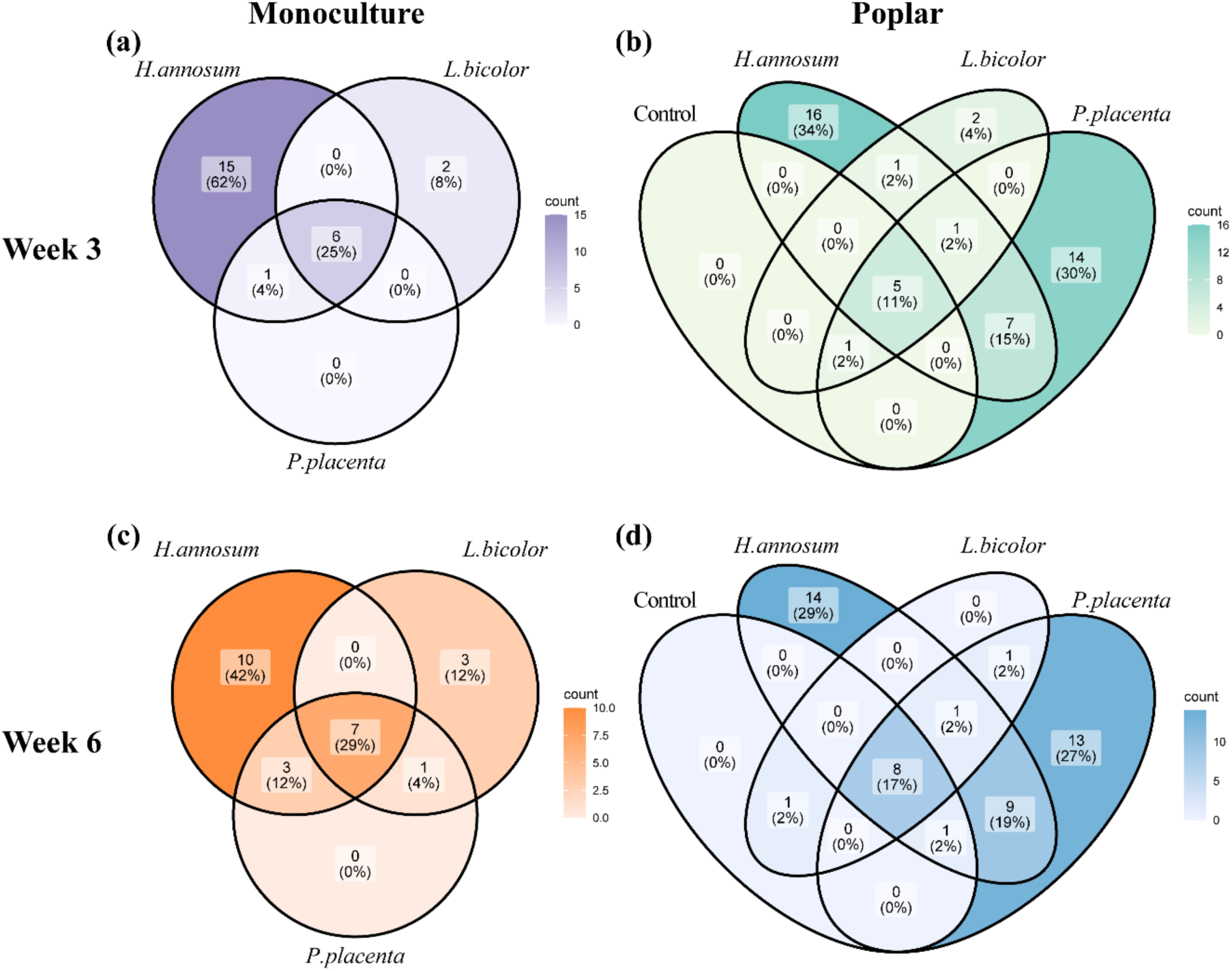
Venn diagrams showing the number and overlap of VOCs detected in belowground from fungal monocultures (*Heterobasidion annosum*, *Laccaria bicolor*, *Postia placenta*) (a, c) and poplar–fungal systems (b, d) at Week 3 (a, b) and Week 6 (c, d) of co-cultivation. Numbers indicate compound counts; percentages represent proportions relative to each system. Color intensity corresponds to the number of VOCs per category.

**Supplementary Figure 3.**
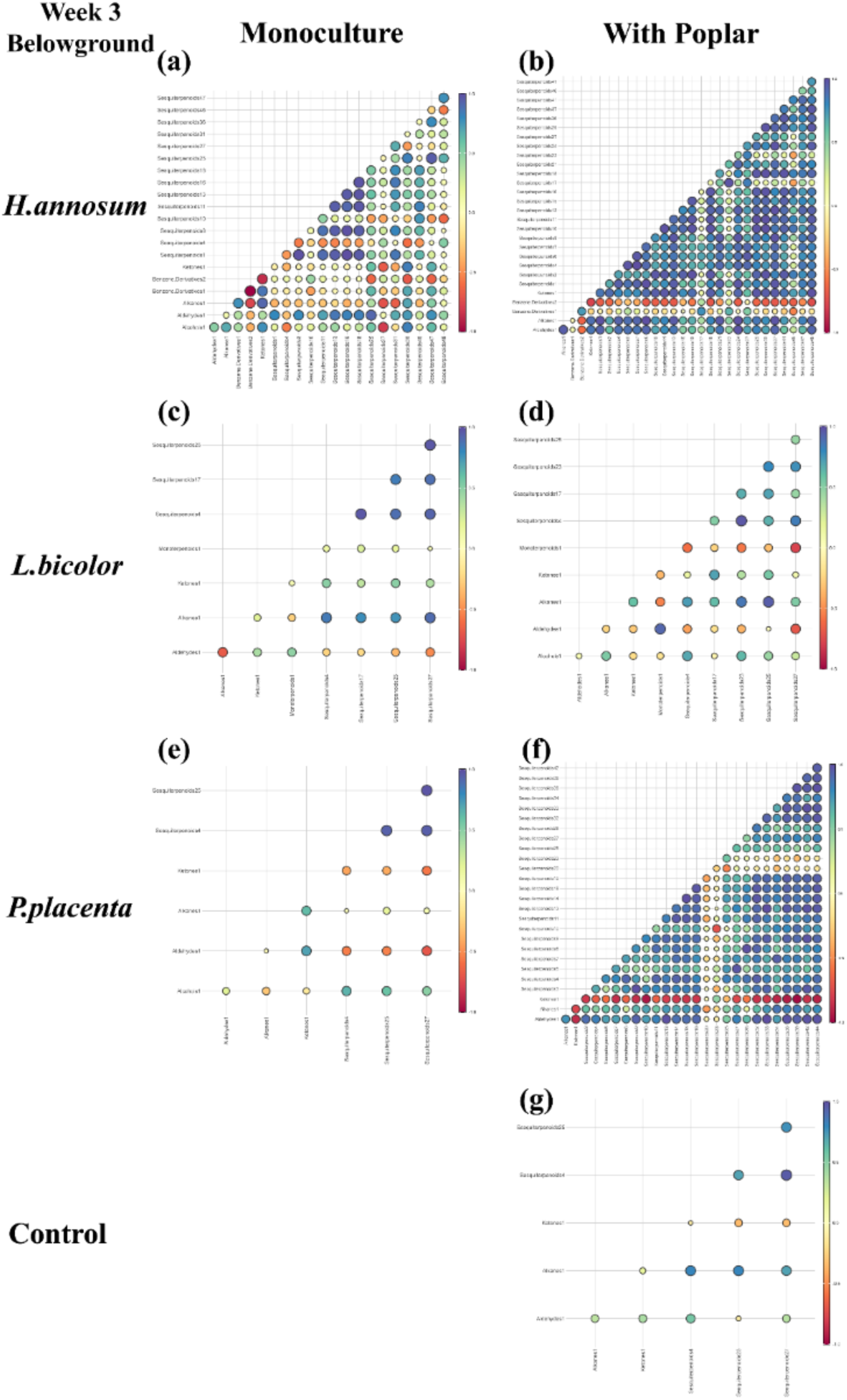
Pairwise Spearman correlation matrices of belowground VOCs detected in fungal monocultures (left) and fungus–poplar systems (right) at Week 3 of co-cultivation. Each subplot shows the strength and direction of VOC–VOC correlations, with circle size and color indicating correlation coefficients (ρ). Only VOCs detected in at least three replicates were included. Panels (a–b) show results for the pathogenic fungus *Heterobasidion annosum*, (c–d) for the ectomycorrhizal symbiont *Laccaria bicolor*, (e–f) for the saprotroph *Postia placenta*, and (g) the poplar control without fungal VOC exposure.

**Supplementary Figure 4.**
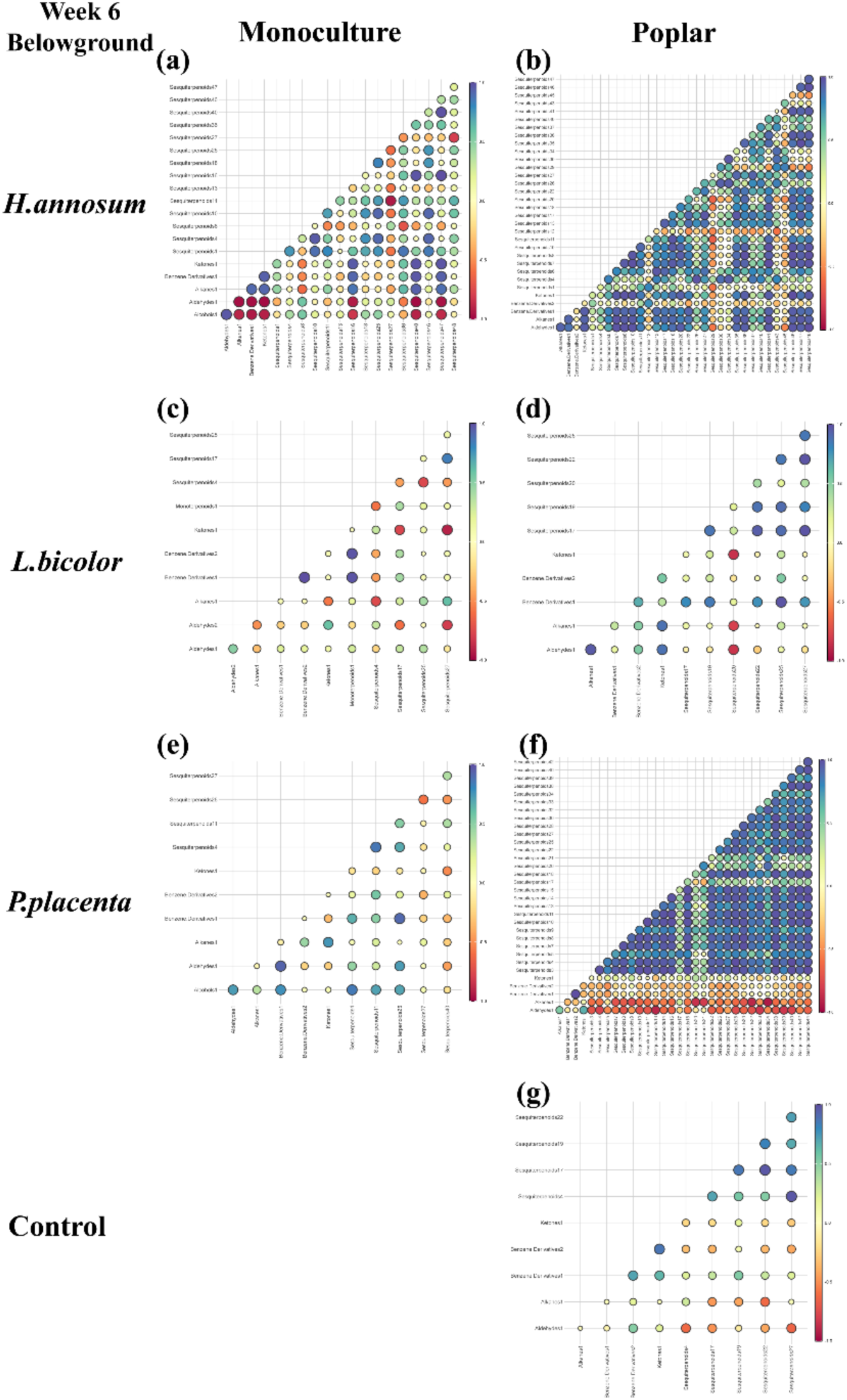
Pairwise Spearman correlation matrices of belowground VOCs detected in fungal monocultures and poplar-fungus systems at Week 6 of co-cultivation. Panels follow the same structure as Supplementary Figure 3, displaying VOC–VOC correlations in (a– b) *Heterobasidion annosum*, (c–d) *Laccaria bicolor*, (e–f) *Postia placenta*, and (g) the poplar control without fungal VOC exposure. Left panels show monocultures; right panels show fungal systems co-cultivated with poplar. Circle size and color reflect correlation strength (Spearman’s ρ).

**Table S1.**
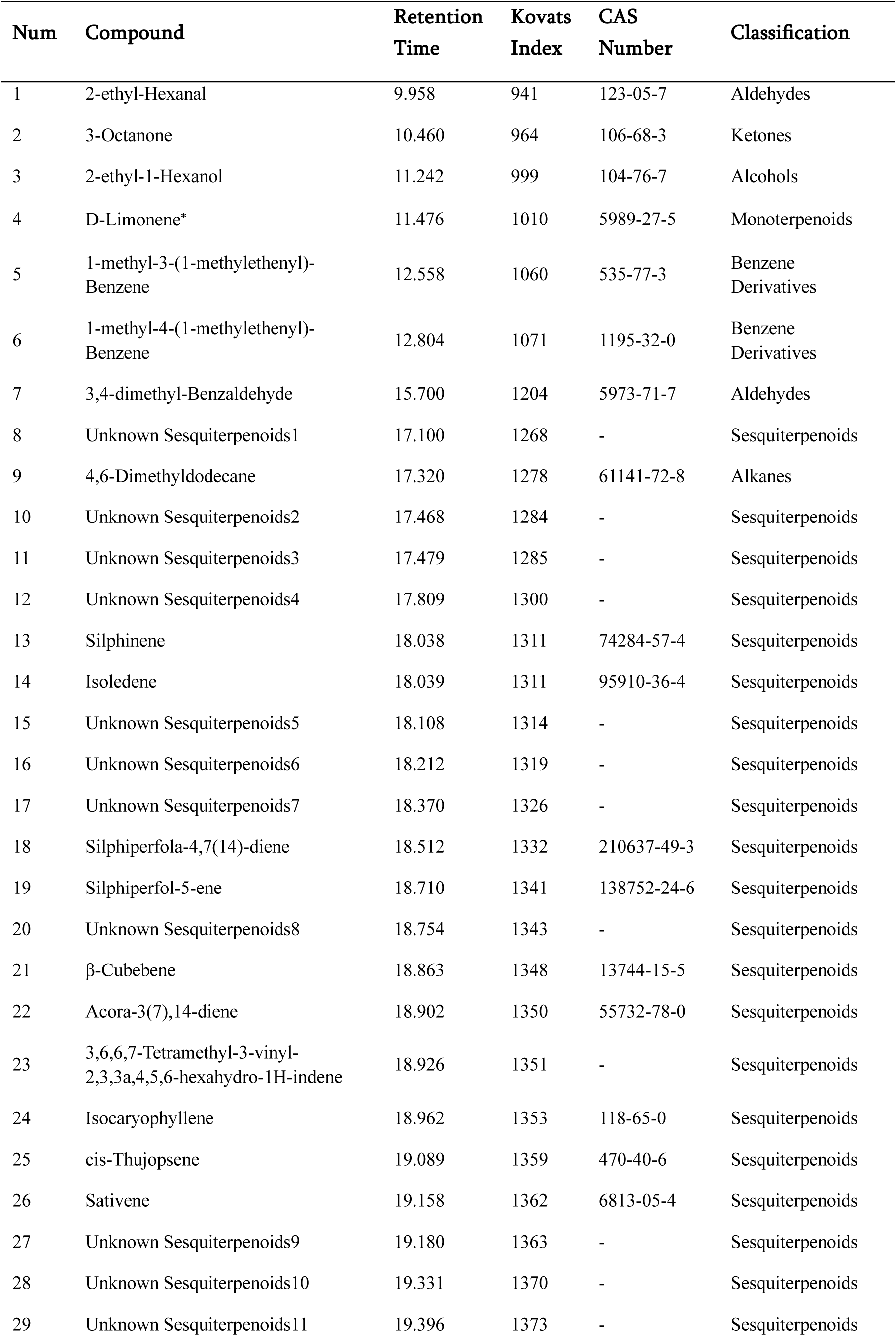
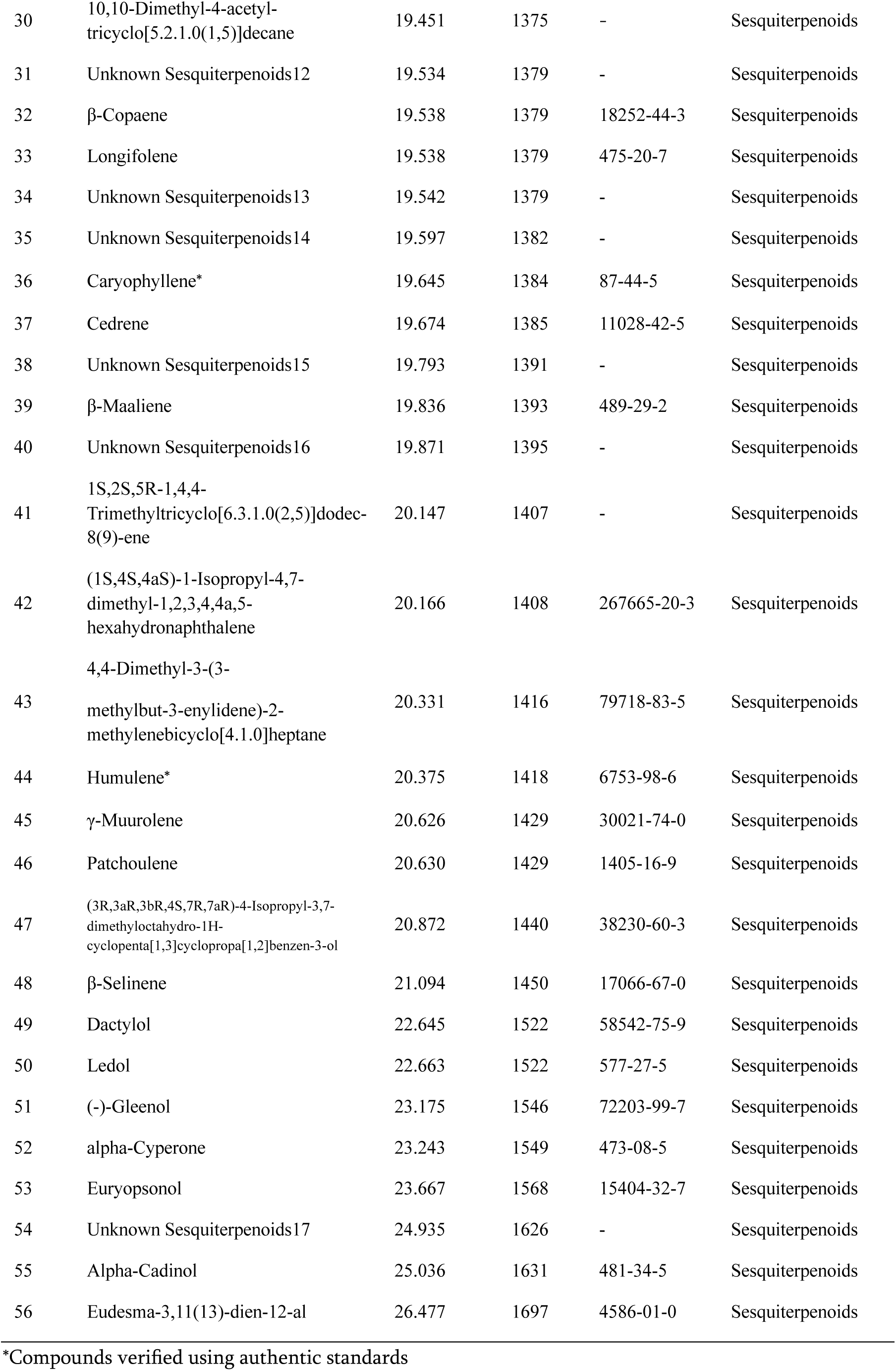
List of Compounds identified in belowground part with their retention times and Kovats index.

**Table S2.**
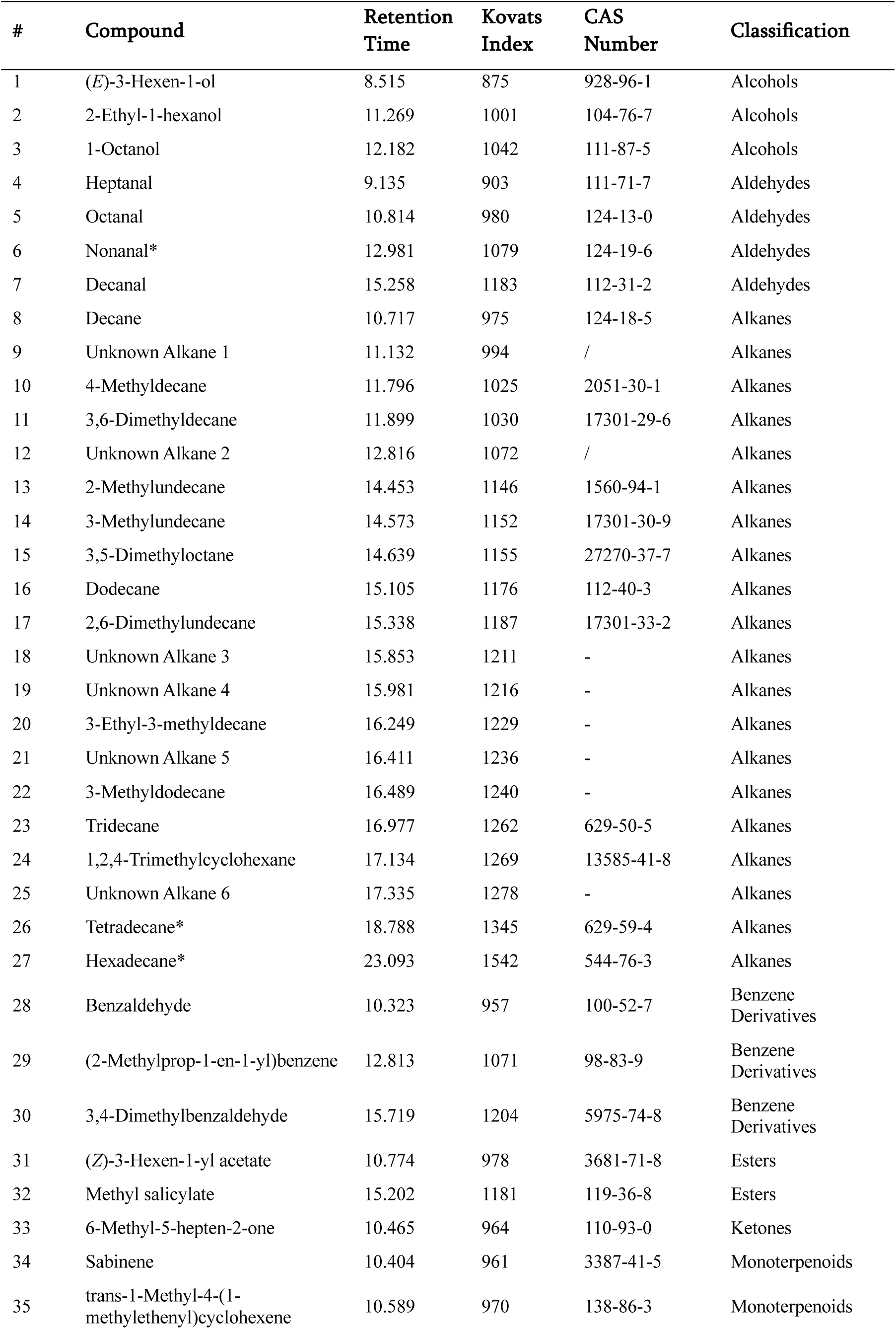
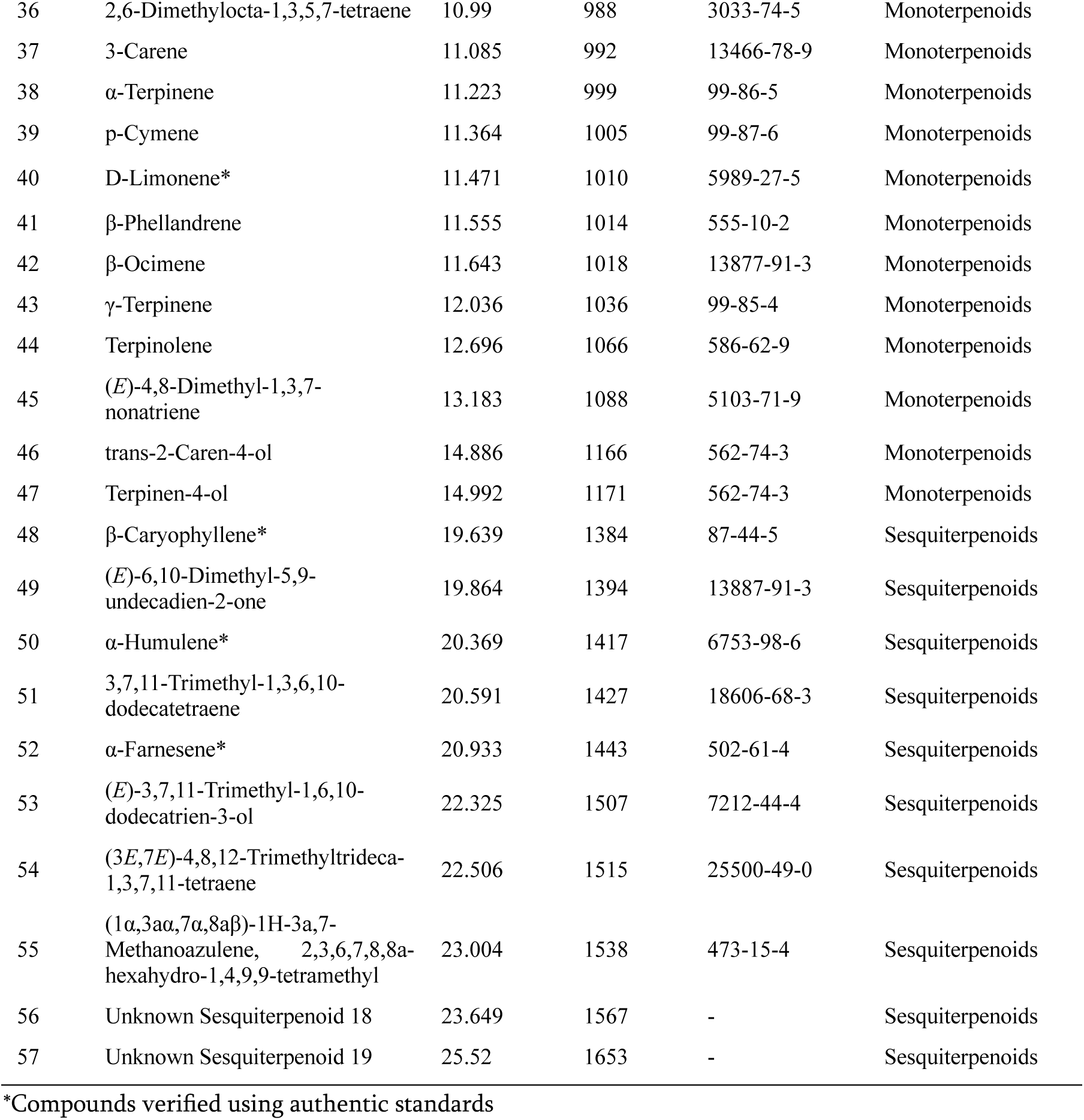
List of compounds identified in aboveground part with their retention times and Kovats index.

**Table S3.**
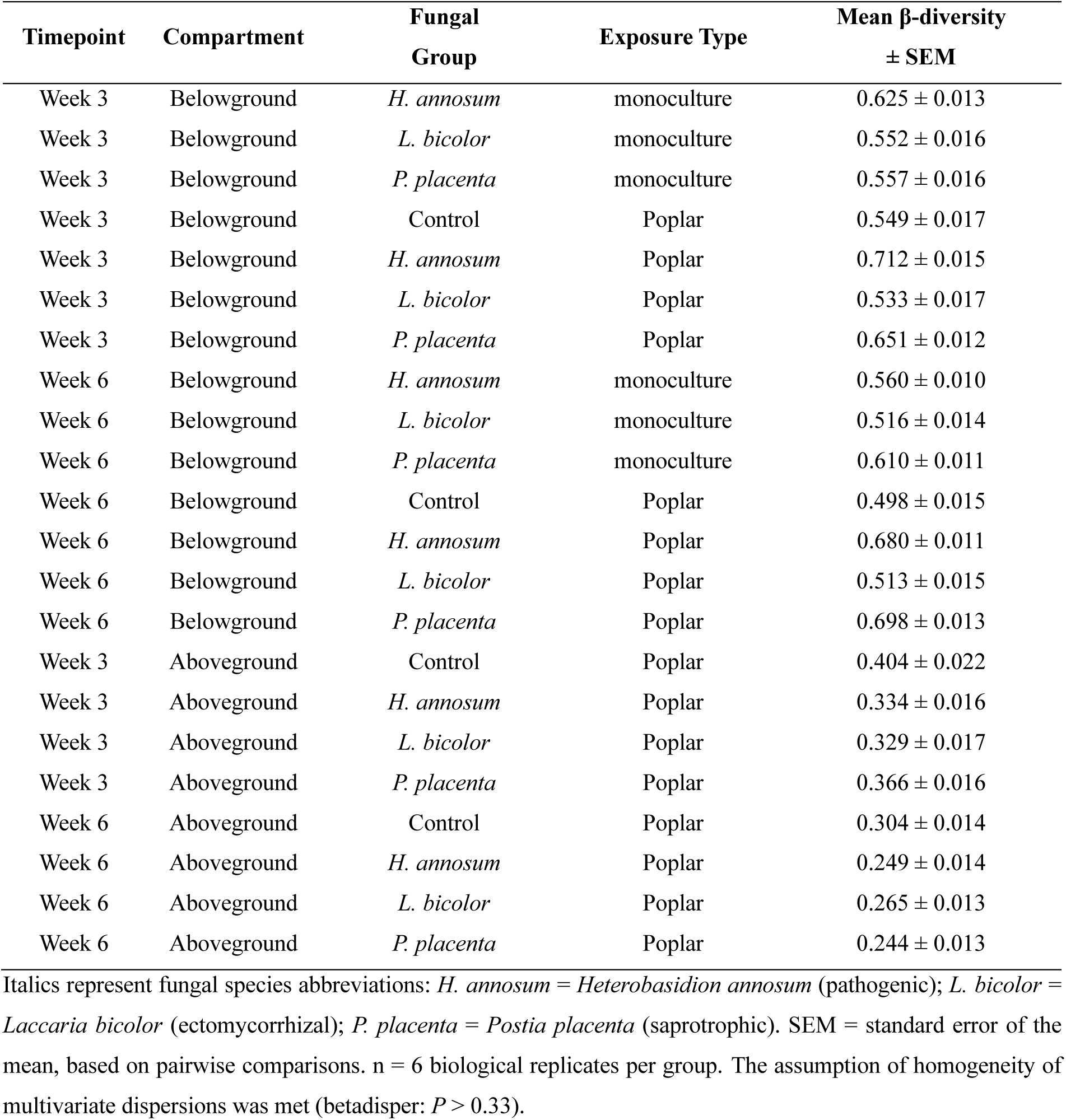
Mean between-group β-diversity (± SEM) in belowground and aboveground VOC communities at weeks 3 and 6 of co-cultivation.

| Timepoint | Compartment | Fungal Group | Exposure Type | Mean β-diversity ± SEM |
| --- | --- | --- | --- | --- |
| Week 3 | Belowground | *H. annosum* | monoculture | 0.625 ± 0.013 |
| Week 3 | Belowground | *L. bicolor* | monoculture | 0.552 ± 0.016 |
| Week 3 | Belowground | *P. placenta* | monoculture | 0.557 ± 0.016 |
| Week 3 | Belowground | Control | Poplar | 0.549 ± 0.017 |
| Week 3 | Belowground | *H. annosum* | Poplar | 0.712 ± 0.015 |
| Week 3 | Belowground | *L. bicolor* | Poplar | 0.533 ± 0.017 |
| Week 3 | Belowground | *P. placenta* | Poplar | 0.651 ± 0.012 |
| Week 6 | Belowground | *H. annosum* | monoculture | 0.560 ± 0.010 |
| Week 6 | Belowground | *L. bicolor* | monoculture | 0.516 ± 0.014 |
| Week 6 | Belowground | *P. placenta* | monoculture | 0.610 ± 0.011 |
| Week 6 | Belowground | Control | Poplar | 0.498 ± 0.015 |
| Week 6 | Belowground | *H. annosum* | Poplar | 0.680 ± 0.011 |
| Week 6 | Belowground | *L. bicolor* | Poplar | 0.513 ± 0.015 |
| Week 6 | Belowground | *P. placenta* | Poplar | 0.698 ± 0.013 |
| Week 3 | Aboveground | Control | Poplar | 0.404 ± 0.022 |
| Week 3 | Aboveground | *H. annosum* | Poplar | 0.334 ± 0.016 |
| Week 3 | Aboveground | *L. bicolor* | Poplar | 0.329 ± 0.017 |
| Week 3 | Aboveground | *P. placenta* | Poplar | 0.366 ± 0.016 |
| Week 6 | Aboveground | Control | Poplar | 0.304 ± 0.014 |
| Week 6 | Aboveground | *H. annosum* | Poplar | 0.249 ± 0.014 |
| Week 6 | Aboveground | *L. bicolor* | Poplar | 0.265 ± 0.013 |
| Week 6 | Aboveground | *P. placenta* | Poplar | 0.244 ± 0.013 |

Italics represent fungal species abbreviations: *H. annosum* = *Heterobasidion annosum* (pathogenic); *L. bicolor* = *Laccaria bicolor* (ectomycorrhizal); *P. placenta* = *Postia placenta* (saprotrophic). SEM = standard error of the mean, based on pairwise comparisons. n = 6 biological replicates per group. The assumption of homogeneity of multivariate dispersions was met (betadisper: $P > 0.33$ ).

